# Dehydrated *Caenorhabditis elegans* stocks are resistant to multiple freeze-thaw cycles

**DOI:** 10.1101/2020.07.27.223560

**Authors:** Patrick D. McClanahan, Richard J. McCloskey, Melanie Ng Tung Hing, David M. Raizen, Christopher Fang-Yen

## Abstract

Ultracold preservation is widely used for storage of genetic stocks of *Caenorhabditis elegans*. Current cryopreservation protocols are vulnerable to refrigeration failures, which can result in the loss of stock viability due to damage during re-freezing. Here we present a method for preserving worms in a dehydrated and frozen form that retains viability after multiple freeze/thaw cycles. After dehydration in the presence of trehalose or glycerol, *C. elegans* stocks can be frozen and thawed multiple times while maintaining viability. While both dauer and non-dauer larvae survive desiccation and freezing, the dauer defective mutant *daf-16* does not survive desiccation. Our technique is useful for storing stocks in a manner robust to freezer failures, and potentially for shipping strains between laboratories.

## INTRODUCTION

Cryopreservation enables collections of biological specimens to be stored for extended periods with little maintenance. Organisms capable of maintaining viability after recovery from long-term cryopreservation include bacterial strains, eukaryotic cell lines, embryos, and small animals such as the genetic model organism *Caenorhabditis elegans* (Brenner, 1974). However, during the freezing process, the specimen may be damaged directly by the formation of ice crystals and by osmotic stress and/or toxicity resulting from the increased concentration of aqueous solutions left as ice forms (Mazur, Leibo and Chu, 1972; Muldrew and McGann, 1990; Muldrew and Mcgann, 1994; Pegg, 2009). For this reason, cryopreservation methods rely on manipulations that reduce the damaging effects of ice crystals.

Most contemporary cryopreservation protocols, including those for *C. elegans* (Brenner, 1974; Stiernagle, 2006), minimize freezing damage by slowly chilling the specimen in a buffer containing a cryoprotective agent. As the specimen is cooled, ice forms in the freezing buffer, concentrating the cryoprotectant. The slow rate of cooling allows the specimen to maintain a safe osmotic balance with the surrounding buffer (Muldrew and Mcgann, 1994) while the increasingly concentrated cryoprotective agent inhibits formation of intracellular ice crystals (Morris *et al*., 2006) and may stabilize lipid membranes and proteins (Rudolph and Crowe, 1985; Crowe, Hoekstra and Crowe, 1992; Burkewitz *et al*., 2012; Julca *et al*., 2012).

Standard preservation techniques in *C. elegans* rely on glycerol added to freezing media as a cryoprotectant (Brenner, 1974; Stiernagle, 2006). Glycerol reduces both the number and size of ice crystals formed during freezing (Morris et al., 2006) by displacing and forming hydrogen bonds with water (Dashnau *et al*., 2006). It may also stabilize lipid bilayers, proteins, and other biomolecules (Clegg *et al*., 1982; Carpenter and Crowe, 1988). Nevertheless, fewer than half the animals survive the freezing and thawing process, and a subsequent refreeze, which may occur after a power or equipment failure, can result in complete loss of stock viability (Stiernagle, 2006).

Many small animals, including tardigrades, rotifers, brine shrimp cysts, and some nematodes, can survive periods of near total desiccation (Crowe, Hoekstra and Crowe, 1992; Wharton, Goodall and Marshall, 2003), a state called anhydrobiosis. Because water is responsible for the damaging effects of freezing, anhydrobiotic organisms are also resistant to freezing damage (Holmstrup, Bayley and Ramløv, 2002). Anhydrobiosis has been described in the *C. elegans* dauer, an alternative third larval stage triggered by stress and/or overcrowding (Erkut et al., 2011). Like many anhydrobiotic organisms (Crowe et al., 1984), *C. elegans* dauers produce the disaccharide trehalose (Erkut et al., 2011). The properties that enable trehalose to function as a xeroprotectant (an agent that protects against dehydration damage) bear some resemblance to those of glycerol in cryopreservation and include its ability to stabilize of lipid bilayers as well as proteins (Carpenter and Crowe, 1988; Crowe, Carpenter and Crowe, 1998). Indeed, trehalose can be used as a laboratory cryoprotectant (Bhandal, Hauptmann and Widholm, 1985; De Antoni *et al*., 1989; Beattie *et al*., 1997), and glycerol can be used as a xeroprotectant (Jin, Taylor and Harman, 1996; Puhlev *et al*., 2001). We therefore hypothesized that *C. elegans* stocks dried in the presence of protective agents might be capable of maintaining viability through multiple freeze-thaw cycles.

In this report we demonstrate that the protective agents trehalose or glycerol allow *C. elegans* to retain viability after dehydration, and that stocks dehydrated in this manner are resistant to freeze-thaw damage. We present a simple method for dehydrating and freezing worms that results in stocks that can survive multiple freeze-thaw cycles. This method is practical for the long-term preservation of *C. elegans* genetic stocks in a manner robust to refrigeration failure.

## MATERIALS AND METHODS

### C. elegans strains and maintenance

The following strains were used in our study: Bristol N2 (WT) and CF1038 *daf-16(mu86)*. We cultured *C. elegans* on OP50 *E. coli* food bacteria on standard NGM agar (Stiernagle, 2006) or standard NGM agar with an addition 200 mM NaCl (salt conditioning experiments only) on 6-cm plates. To prepare worms, we picked five young adult hermaphrodites onto each plate and allowed the population to grow at ambient temperature (RT, 16–24 °C) for three weeks prior to conducting experiments. The food bacteria became depleted about one week after picking, so these animals were without food for about two weeks. With the exception of freezing and long term storage experiments, culturing and experiments were performed at RT.

### Buffers and solutions

M9 buffer, S buffer, and soft agar freezing media were made according to standard methods (Stiernagle, 2006). Alternative freezing (AF) media consisted of 15.1 g trehalose, 17.7 ml DMSO, and M9 buffer to a total volume of 500 ml (Kevin O’Connell, personal communication). All cryo/xeroprotectants were prepared immediately prior to use as solutions in M9 at 30% concentration (v/v for glycerol and DMSO, w/v for trehalose).

### Liquid freezing (LF)

The LF procedure was previously described (Stiernagle, 2006) except that starved animals from three week old plates were used instead of freshly starved animals, and M9 buffer was used instead of S buffer (Brenner, 1974). *C. elegans* were washed from growth plates into a 15 mL conical centrifuge tube using 0.7 mL M9 per plate. An equal volume of M9 with 30% (v/v) glycerol was added to the tube and mixed by gentle vortexing to generate “worm freezing mixture”. We added 1 mL of worm freezing mixture into each 1.8 mL cryovial (USA Scientific) and 5 μL was pipetted onto a food plate for counting of the viable worm population prior to freezing (see counting section). The cryovials were capped, placed in a polystyrene foam shipping container (18.5 cm x 18.5 cm x 14 cm outer dimensions with 2.2-2.5 cm wall thickness) to slow cooling, and frozen at approximately −78.5° C in a larger cooler lined on the bottom and two sides with dry ice slabs. After freezing and / or simulated refrigeration failures, vials were placed at RT for 10-15 min, mixed by gentle vortexing, and either 10 μL (1X freeze thaw) or the full 1 mL (simulated refrigeration failures) of the thawed media was pipetted onto a food plate for counting viable worms.

### Soft agar freezing (SA)

The SA procedure was as previously described (Stiernagle, 2006) except that starved animals from three week old plates were used instead of freshly starved animals. Worms were washed from growth plates into a 15 mL centrifuge tube using 0.7 mL S buffer per plate and placed on ice. After 15 min, an equal volume of molten, 50°C soft agar freezing solution was added to the conical and mixed by gentle vortexing. Freezing and recovery was identical to that described for the LF procedure.

### Trehalose-DMSO alternative freezing (AF)

The trehalose-DMSO freezing protocol was similar to one in use elsewhere (Kevin O’Connell, personal communication). Worms starved for 3 weeks were washed from their growth plates with M9, pelleted at 700 rcf in a clinical centrifuge for 3 min, washed once in trehalose-DMSO freezing buffer, and resuspended in trehalose-DMSO freezing buffer to make a worm freezing mixture. We added 1 mL of worm freezing mixture into each cryovial and placed them in a −78.5° C dry ice-lined cooler in a foam shipping container.

### Desiccation with xeroprotectants

Animals were transferred from several plates into a 15 mL centrifuge tube by floating them in M9 buffer using a transfer pipette (USA Scientific), then pelleted (5 min at 700 rcf), washed in 10 mL of fresh M9, pelleted again, and transferred to a 1.5 mL microcentrifuge tube. We added M9 to the tube to adjust the volume to approximately 50 μL per worm plate used. After brief vortexing, this worm mixture was combined with equal parts of 30% (w/v for trehalose, v/v for glycerol and DMSO, or M9 alone) xeroprotectant solution in M9 for a final concentration of 15%. We added 50 μL of well-mixed worm-xeroprotectant mixture to each cryovial and 1 μL was pipetted onto a food plate for baseline counting (see counting section). The uncapped cryovials were placed in an airtight box also containing approximately 5 g anhydrous calcium sulfate desiccant (Hammond Drierite 23001, 8 mesh) per cryovial and allowed to dry for 48 h at RT. Trehalose-containing buffer dried to a soft solid, and glycerol-containing buffer dried to a viscous liquid. See also Supplementary Protocol (File S2).

### Simulated refrigeration failure and freeze-thaw cycling

Cryovials containing *C. elegans* were placed in an expanded polystyrene foam shipping container (same as used for conventional freezing) which was placed in an insulated container lined with dry ice slabs on the bottom and two sides. The following day, cryovials subjected to a single freeze-thaw (1X) were removed, thawed, and rehydrated (see rehydration and recovery). The remaining vials were kept in the shipping container, which was stored at RT for approximately 8 h and then returned to the −80 °C to simulate a single refrigeration failure (2X). The protocol was repeated the next day to simulate a second refrigeration failure (3X). On the last day, cryovials subject to an additional three rounds of freeze thaw cycling (6X) were placed in a cardboard cryostorage box and switched between RT and −80 °C at 1 h intervals.

### Counting animals before and after freezing and/or desiccation

For all counting, we pipetted some or all of the volume of worm suspension onto an NGM plate that had been seeded with a lawn of OP50 bacteria. To minimize underestimation due to worms sticking to the inside of the pipette, we pipetted the worm suspension up and down once to precoat the inside of the pipette tip with worms before transferring worms. We counted animals that moved spontaneously or in response to either tapping the plate or to touching them with a worm pick made of platinum-iridium wire (Tritech PT-9010, 254 μm diameter). Counted animals were removed from the plate.

To estimate the number of worms prior to freezing and/or desiccation, we counted animals in a small sample of the worm suspension (5 μL for SA, LF, and AF, 1 μL for desiccation). Sample volumes were chosen in order to maintain the number of counted worms in roughly the range of 100 - 500. Dauer, L4, and adult animals were counted within 1 h after pipetting the worm sample on the agar surface, and the remaining larvae counted the following day, allowing L1s to reach a more easily visible size. The number of worms placed in each tube for freezing and/or desiccation ranged from 3800 to 59200 (mean 13221).

To count the number of animals surviving after recovery, we used a similar procedure, except counting was performed a few hours after recovery and then repeated daily for six days to allow animals additional time to recover. For LF, SA, and AF experiments, we allowed the vials to thaw for 15 min, vortexed the contents briefly, and pipetted 10 μL (single freeze-thaw samples) or the full 1 mL (one or more simulated refrigeration failures) of the thawed suspension onto a food plate. For desiccation experiments, we allowed the vials to thaw for 10 min (if frozen or chilled), rehydrated the worms in 50 μL of M9 at RT for 10 min, mixed the worm suspension by tapping the side of the cryovial, and pipetted 5 – 6.5 uL (WT in trehalose or glycerol buffer except 1 month storage at 20°C) or the full 50 uL (all *daf-16*, WT in plain M9, WT in DMSO buffer, and WT in trehalose or glycerol buffer stored for 1 or 3.5 month at 20°C) of the worm suspension onto a food plate.

Since some worms took several days to recover, and plates were assayed once per day, there was some uncertainly in the identification of developmental stages. During counting, we classified surviving worms as L1-L3 larvae, dauer larvae, or L4 / adult. Because dauers develop into L4s, recovered L1-L3 larvae were assumed to have survived freezing and/or drying as non-dauer larvae. Therefore, L1-L3 survival rate was estimated by dividing the number of L1-L3 larvae recovered by the number of L1-L3 larvae desiccated / frozen. Similarly, because the food-rich, low population density recovery plate conditions favor dauer exit rather than dauer entry, recovered dauers were assumed to have survived as dauers as opposed to having entered the dauer state after recovery. Therefore, dauer survival rate was estimated by dividing the number of dauers recovered by the number of dauers desiccated / frozen. In contrast, recovered L4 / adults could have developed from surviving dauer larvae or L1-L3 larvae, or survived as L4 / adults. Therefore, these animals were not used in estimations of survival by type, but did contribute to calculations of overall survival. The type-specific survival rates may be underestimated because some surviving L1-L3s and dauers could have developed into L4s or adults after recovery but before being counted.

### Imaging dehydration and rehydration of worms in a single droplet

To capture images of *C. elegans* during dehydration and rehydration, we prepared 3-week starved N2 animals in M9 solution containing 15% trehalose, then pipetted a 10 μl droplet of the suspension onto a polydimethylsiloxane (PDMS, Dow Corning Sylgard 184) surface. We used PDMS for its optical clarity and non-stick nature, enabling us to easily remove the dehydrated flake. We placed the PDMS layer in a clear polystyrene dish, along with anhydrous glycerol as a desiccant, on the stage of a Leica M165 FC stereo microscope. Images were captured at 1 frame per minute using a 5 MP Imaging Source CMOS camera. The next day, we transferred the sample to an OP50-seeded plate using forceps, covered it with 10 μl water, and recordied images on the stereo microscope as animals recovered.

### Plotting and statistics

All statistical tests and plotting were performed in MATLAB. All bar graphs show mean ± standard error of the mean. The number of replicates for each experiment is given in the figure captions or text. Circles represent data from individual replicates. We used α = 0.05 for determining statistical significance. Statistical tests and results are described in the results.

## RESULTS

### C. elegans frozen by established methods do not survive repeated freeze-thaw cycles

It has been reported that a power failure can result in loss of all *C. elegans* stocks stored in a −80° C freezer (Stiernagle, 2006). To confirm this observation, and to obtain baseline data against which to compare subsequent results, we quantified the viability of *C. elegans* frozen by standard methods after single or repeated freeze-thaw cycles.

Two *C. elegans* freezing protocols are in common use (Stiernagle, 2006). In liquid freezing (LF), starved worms are frozen in liquid buffer supplemented with 15% glycerol as cryoprotectant. In soft agar freezing (SA), a small concentration of low-melting temperature agar is added to the freezing medium; the agar keeps worms suspended throughout the medium, such that repeated small portions of the frozen medium can be removed and thawed instead of thawing the entire tube. We froze WT animals using both the LF and SA protocols, then measured their survival rates upon thawing with or without one or two simulated refrigeration failures. In a simulated refrigeration failure, tubes housed in an insulated expanded polystyrene box were thawed at RT for 8 h (see Methods) and then refrozen.

Survival after a single freeze and thaw, corresponding to the normal procedure for cryopreservation, was 36 ± 4% for LF and 41 ± 11% for SA **(Fig. 1a-b)**, consistent with the previously reported range of 35-45% (Stiernagle, 2006). After a second freeze-thaw cycle, survival dropped precipitously to 0.037 ± 0.032% for LF and 0.020 ± .001% for SA, although a few individuals from each of the three vials tested per condition survived. After three freeze-thaw cycles, there were no survivors out of over 200,000 worms frozen. Interestingly, dauer larvae, which are resistant to a wide range of stressors including cold (Savory, Sait and Hope, 2011), survived at a rate similar to non-dauer larvae, confirming prior observations that dauer larvae are not more resistant to freezing than non-dauer larvae (Stiernagle, 2006).

**Figure 1.**
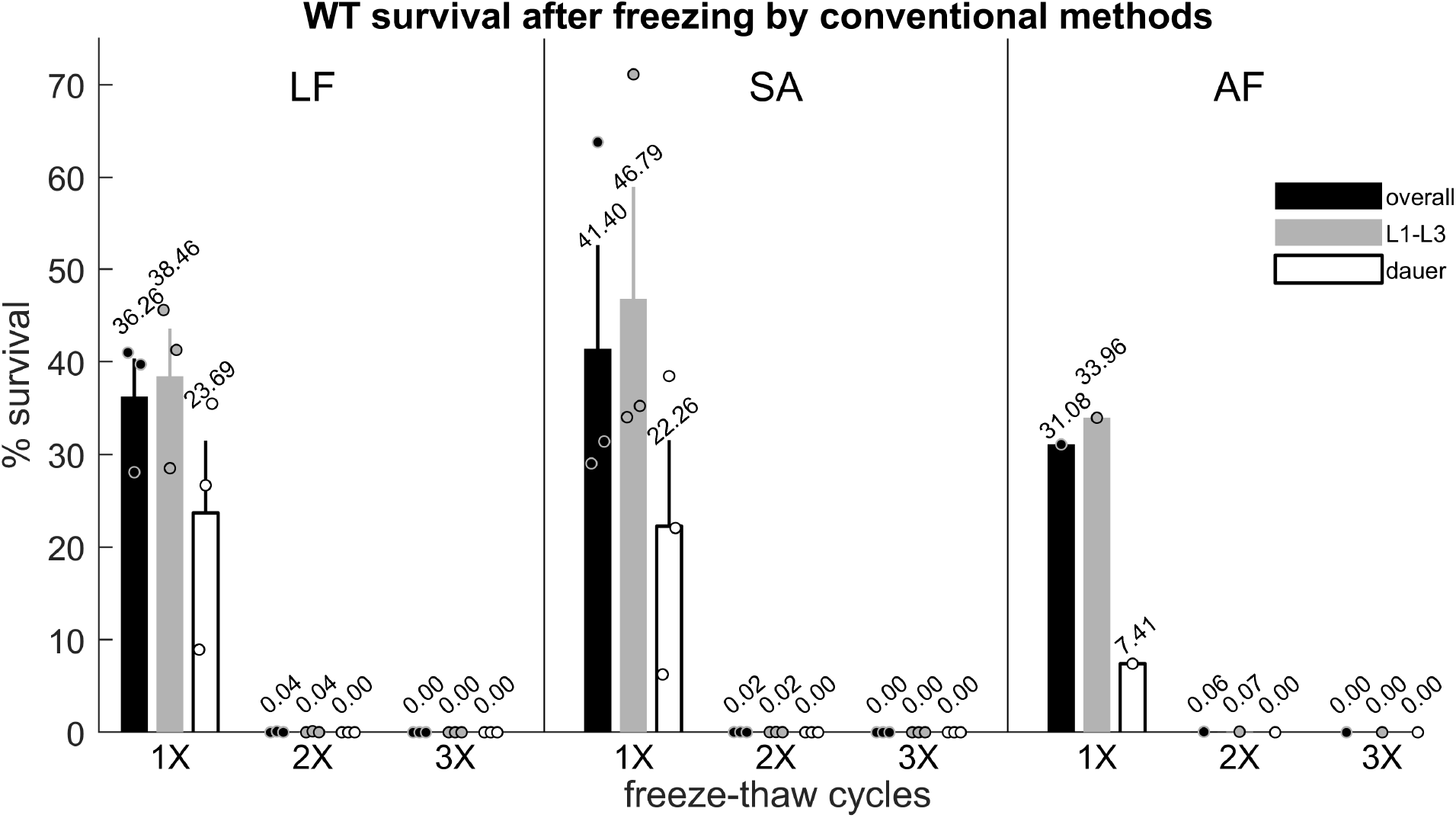
Worms frozen using conventional media lose viability after repeated freeze-thaw cycles. **(Left)** Survival of worms frozen in liquid freezing (LF) media (15% glycerol in M9), three replicates per condition. **(Center)** Survival of worms frozen in soft agar (SA), three replicates per condition. **(Right)** Survival of worms frozen in alternative freezing (AF) buffer, one replicate per condition. Black bars indicate overall survival. Dark gray and unfilled bars indicate L1-L3 and dauer survival, respectively. Error bars represent ± SEM. Circles represent survival in individual replicates. Each replicate contains between 23,200 and 59,200 worms. No living animals were recovered after three freeze-thaw cycles using these methods.

While conventional freezing buffers include glycerol as a cryoprotectant, some laboratories have developed *C. elegans* freezing buffers with alternative cryoprotectants, for example the disaccharide trehalose (Mitani, 2009). To determine if a different type of cryoprotectant might better protect *C. elegans* over multiple freeze thaw cycles, we also tested an alternative freezing (AF) buffer including both trehalose and DMSO (see Methods). This buffer performed similarly to LF and SA **(Fig. 1c)**.

Together, these results confirm that standard frozen *C. elegans* stocks are highly vulnerable to freeze-thaw cycles.

### Exogenous trehalose or glycerol can serve as C. elegans xeroprotectants

Since damage during freezing is thought to be partly caused by the formation of ice crystals (Mazur, 1963; Pegg, 2009), we hypothesized that desiccation prior to freezing might improve *C. elegans* survival after repeated freeze-thaw cycles. However, like freezing, desiccation also causes damage to *C. elegans* (Ohba and Ishibashi, 1981; Gal, Glazer and Koltai, 2004; Erkut *et al*., 2011). We therefore first sought to develop a simple method for dehydrating worms while preserving viability.

Some compounds, such as trehalose and glycerol, can serve as both cryoprotectants and xeroprotectants (Julca *et al*., 2012). *C. elegans* adaptation to osmotic stress involves the production of endogenous glycerol (Lamitina *et al*., 2004), and the ability of dauer larvae to survive desiccation depends on genes required for production of trehalose (Erkut *et al*., 2011).

We asked whether the presence of exogenous protective agents might improve desiccation tolerance in various developmental stages of *C. elegans*. In addition to trehalose and glycerol, we also tested dimethylsulfoxide (DMSO), which has been used as a cryoprotectant (Pegg, 2009), including for the nematodes *C. briggsae* (Hwang, 1970) and *C. elegans* (Kevin O’Connell, personal communication). We desiccated starved *C. elegans* in droplets of M9 buffer alone or M9 supplemented with 15% trehalose **(Fig. 2, Supplemental Video File S1)**, 15% glycerol, or 15% DMSO for 48 h in uncapped cryotubes placed in sealed containers containing a desiccant. We then rehydrated and transferred the desiccated worms to the surface of NGM plates seeded with bacteria **(Fig. 2)** and tracked their recovery.

**Figure 2.**
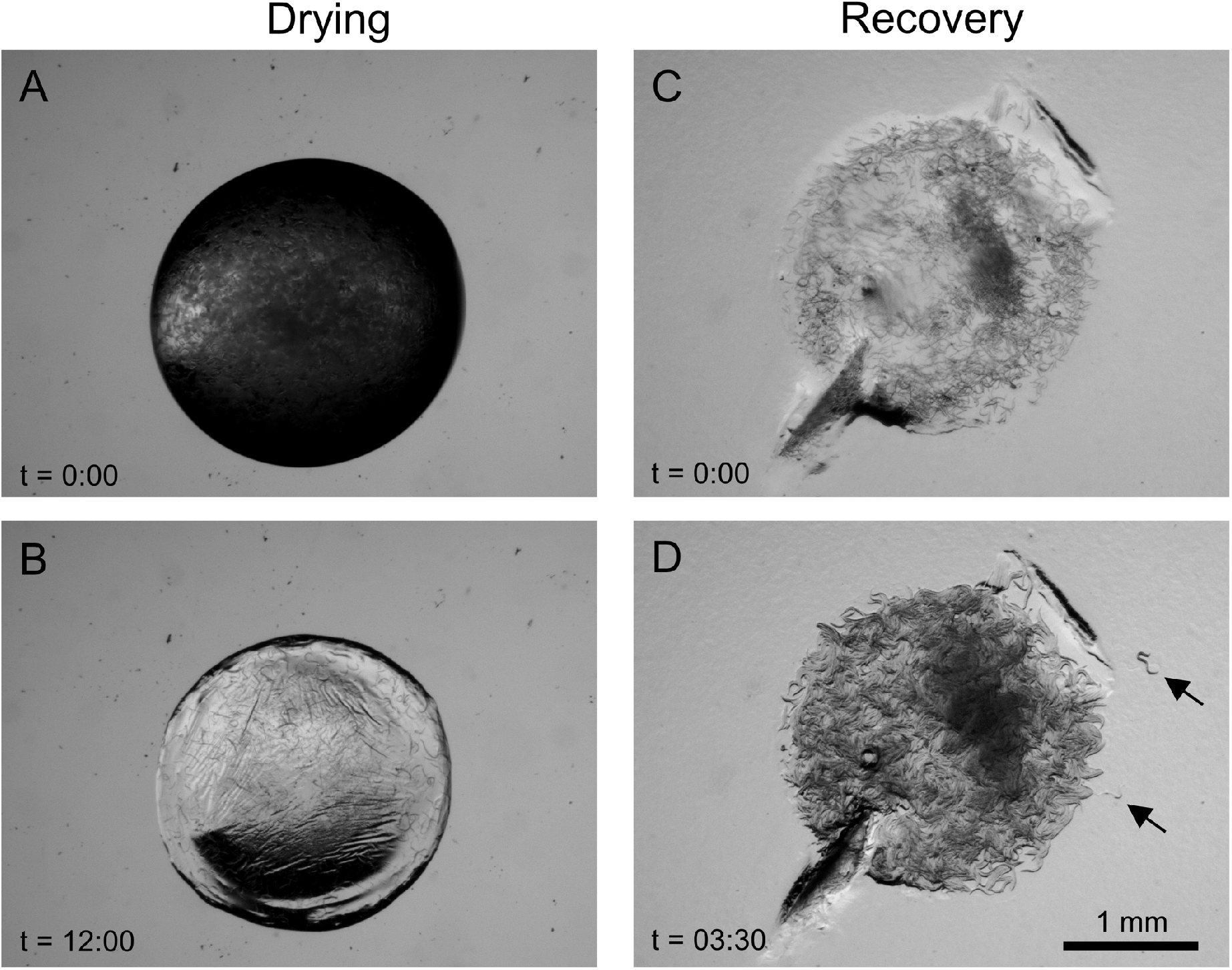
Drying and recovery of *C. elegans*. (A) Worms in a droplet containing trehalose. (B) After 12 h of dehydration, the droplet has dried to a solid. (C) The same sample after being transferred to an OP50-seeded plate. (D) The same sample 3 h, 30 min later. Several animals (arrows) have moved away from the drop. See also Supplementary Video File S1.

Addition of either trehalose or glycerol led to a robust increase in survival after desiccation: overall survival was 5.9 ± 1.2% for worms dried in trehalose media and 3.3 ± 1.0% for worms dried in glycerol media. In contrast, 0.16 ± 0.05% animals survived drying in M9 alone, and none survived in media containing DMSO **(Fig. 3)**, perhaps due to the toxicity of DMSO at high concentrations (Ura *et al*., 2002).

**Figure 3.**
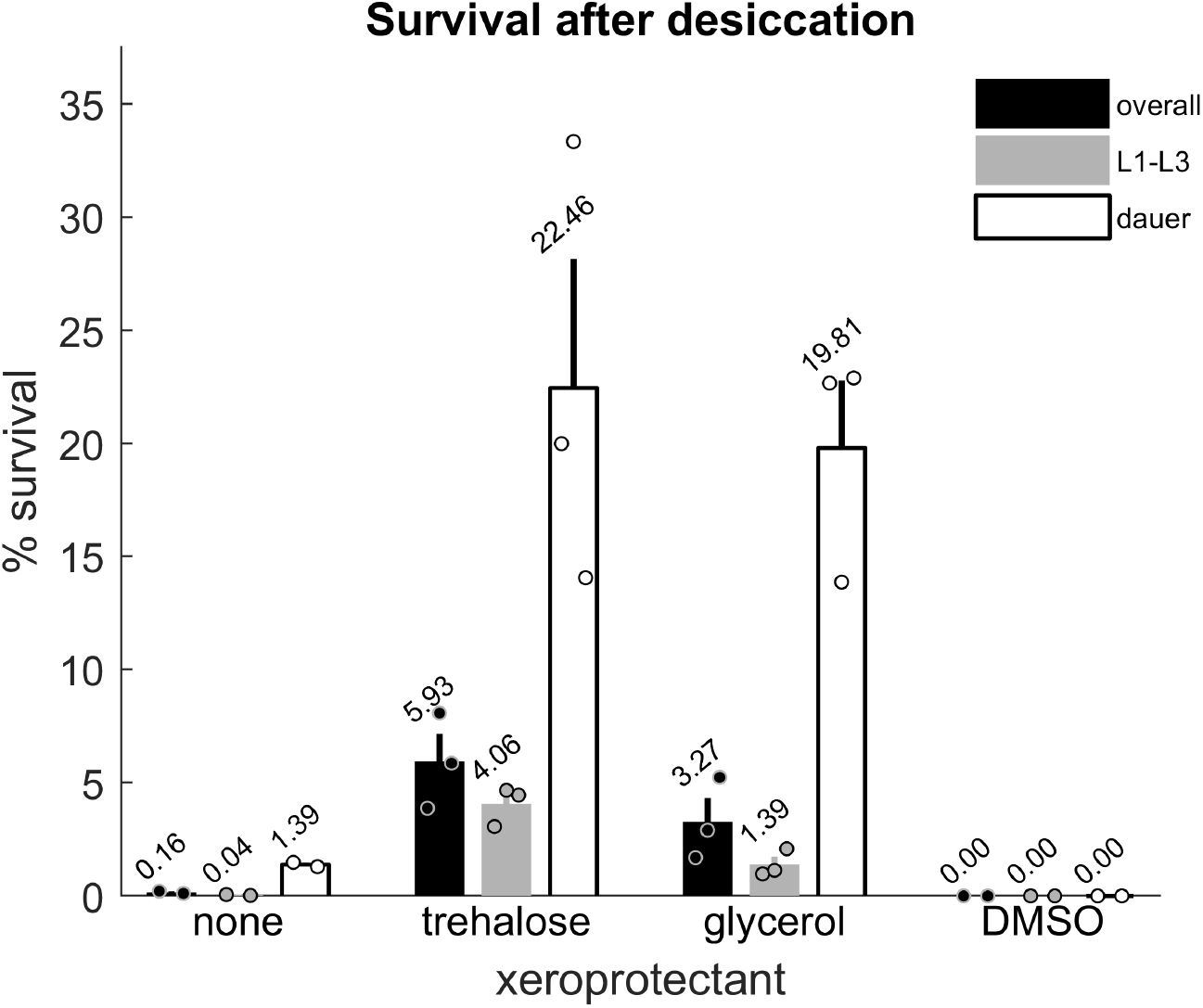
Xeroprotectants improve survival after desiccation. Survival of worms dried in M9 buffer alone or M9 buffer supplemented with 15% trehalose or with 15% glycerol. There are two replicates for M9 alone and three replicates per condition for M9 + xeroprotectant. Each replicate contains approximately 7000 - 22500 animals. Black bars indicate overall survival. Dark gray and light gray bars indicate L1-L3 and dauer survival, respectively. Error bars represent ± SEM. Circles represent individual replicates.

In the absence of added xeroprotectants, dauer larvae have been reported to survive desiccation better than non-dauer larvae (Ohba and Ishibashi, 1981; Erkut *et al*., 2011). We reproduced this observation (Fig. 2). We then asked whether survival after desiccation in the presence of trehalose or glycerol is similarly higher for dauer animals. For ease in counting, we grouped worms into two categories: dauer larvae or L1-L3 larvae. Surviving L4s and adults were excluded because they could have survived desiccation as either dauer or non-dauer larvae and molted prior to being counted. With either trehalose or glycerol, dauer survival was increased several fold. Survival of non-dauer larvae was also increased, albeit more modestly, **(Fig. 3)**.

These results show that exogenous trehalose and glycerol can serve as xeroprotectants for both dauer and non-dauer larvae.

### Desiccated C. elegans stocks withstand multiple freeze thaw cycles with little reduction in viability

Having established that exogenous trehalose and glycerol promote tolerance to desiccation, we next asked whether *C. elegans* stocks desiccated in the presence of these agents can maintain viability through freeze-thaw cycles. We subjected worms dried with exogenous trehalose and glycerol to one, two, or three freeze-thaw cycles, as before. We then subjected some samples to an additional three rapid freeze-thaw cycles for a total of six freeze-thaw cycles before rehydration and recovery.

We observed no significant correlation between number of freeze thaw cycles and overall survival for samples dried in trehalose buffer (coefficient of correlation *ρ* = −0.079 and p = 0.78) or glycerol buffer (*ρ* = −0.23 and p = 0.41) **(Fig. 4)**, indicating that dehydrated stocks of *C. elegans* do not lose viability when repeatedly frozen and thawed up to at least six cycles.

**Figure 4.**
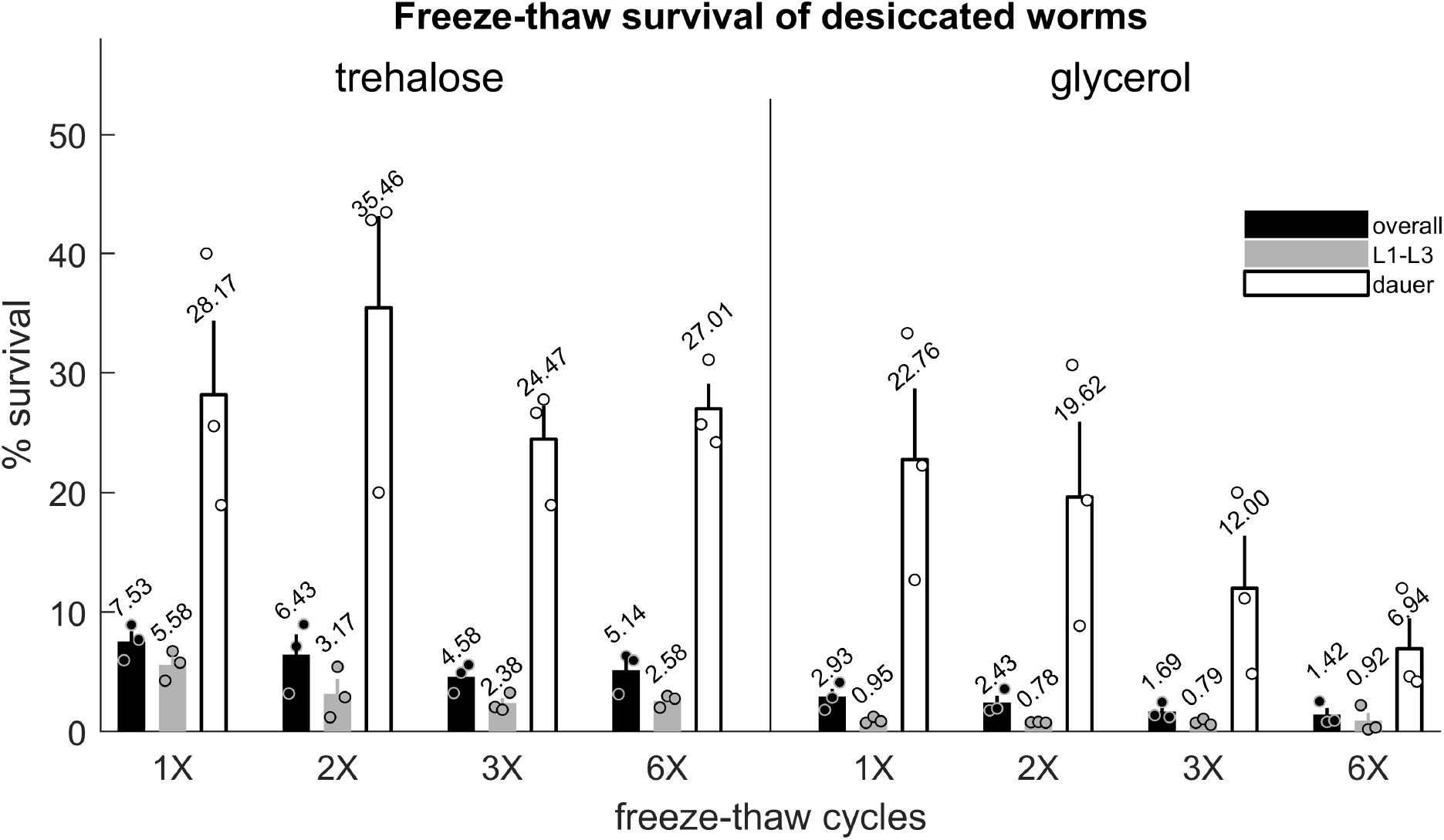
Desiccated worms survive multiple freeze-thaw cycles. Survival of worms dried in M9 buffer supplemented with 15% trehalose or 15% glycerol and subjected to one, two, three, or six freeze-thaw cycles. There are three replicates per condition. Black bars indicate overall survival. Dark gray and light gray bars indicate L1-L3 and dauer survival, respectively. Error bars represent ± SEM. Circles represent individual replicates. Each replicate contains approximately n = 15000 - 22500 animals.

These results show that *C. elegans* desiccated with exogenous trehalose and glycerol are robust to multiple freeze-thaw cycles.

### Desiccated C. elegans retain viability indefinitely at low temperature

The ability to survive multiple freeze-thaw cycles represents a major advantage of desiccated *C. elegans* stocks over conventional *C. elegans* freezing protocols. To further evaluate the suitability of our method for long term storage, we asked how long *C. elegans* dehydrated with a protective agent remain viable at various temperatures. We dehydrated worms in trehalose or glycerol media and stored sealed tubes at either −80° C, −20° C, +4° C, or +20°C. We then assayed survival after one day, one week, one month, and 3.5 months **(Fig. 5)**.

By fitting with an exponential function s(t) = s_0_ exp(−t/*τ*) we estimated the characteristic decay time *τ* for overall survival at each temperature. As expected, survival decayed more rapidly with increased temperature. Estimated decay times were *τ* = 267 d (95% confidence interval 82-∞ d) at −20° C, *τ* = 61 d (95% CI 18-∞ d) at +4° C, and *τ* = 3.2 d (95% CI 3,0-3.4 d) at +20° C for worms dried in trehalose buffer, and *τ* = 51 d (95% CI 24-∞ d) at −20° C, *τ* = 50 d (95% CI 28-237 d) at +4° C, and *τ* = 3.0 d (95% CI 2.9-3.1 d) at +20° C for worms dried in glycerol buffer.

**Figure 5.**
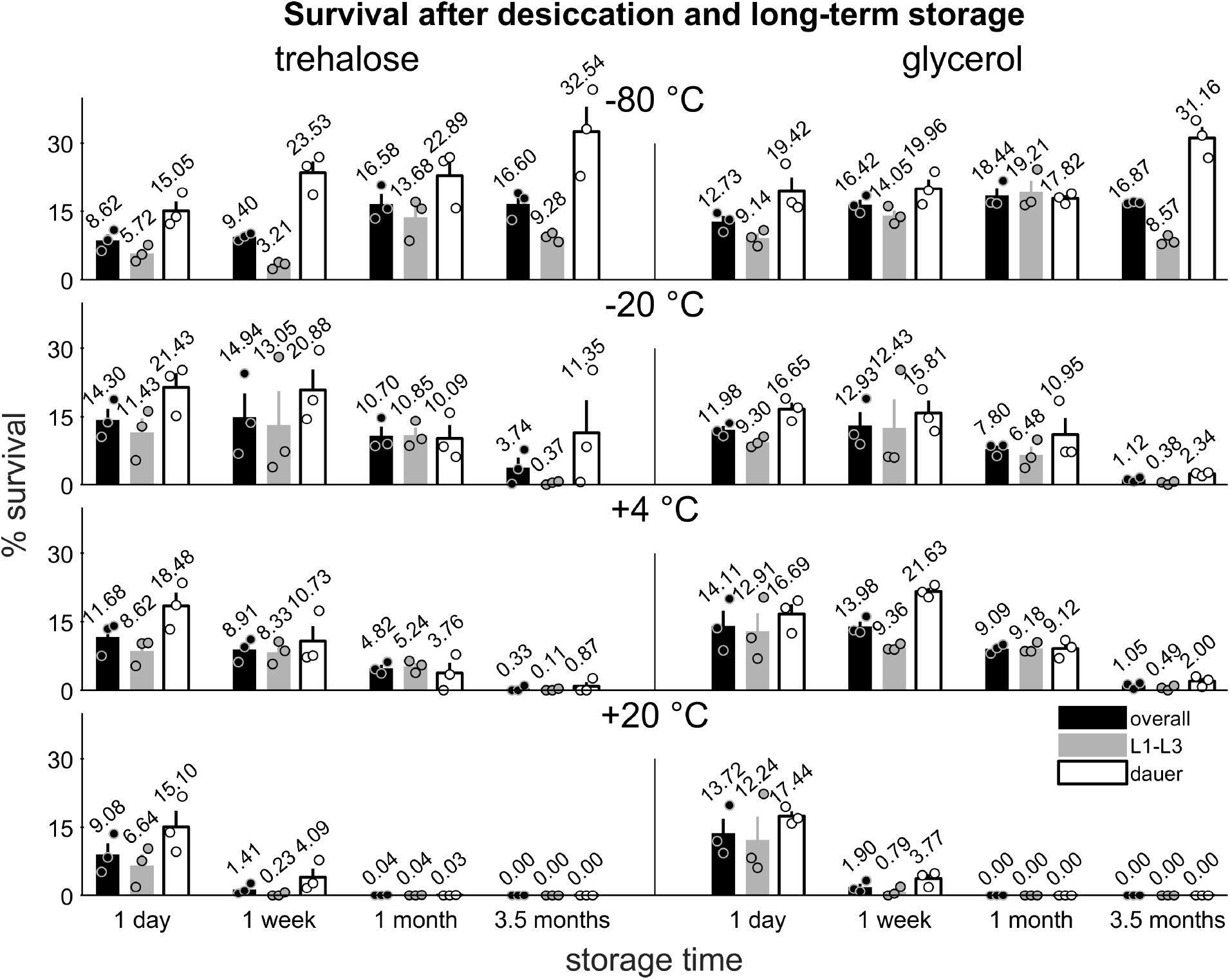
Long term survival of desiccated stocks at various temperatures. Survival of wild type *C. elegans* desiccated in M9 supplemented with trehalose (left) or glycerol (right) and stored for one day, one week, one month, or 3.5 months at −80, −20, 4 or 20 °C (top to bottom). Overall survival is indicated by black bars, L1-L3 larval survival is indicated by dark gray bars, and dauer survival is represented by light grey bars. Error bars represent ± SEM. Circles represent individual replicates. There are three replicates per condition, with approximately n = 3800 – 7800 animals per replicate.

We noted that the proportion of dauers surviving in this set of experiments was lower than in previous experiments. This may have been due to fluctuations in cultivation temperature which ranged from 16-24 °C (see Methods).

These results suggest that desiccated stocks can be stored indefinitely under ultracold temperatures, but also remain viable for extended periods under ordinary refrigeration and several days at room temperature. These findings may be useful for managing storage of desiccated stocks after a freezer failure.

### Starvation time affects survival after desiccation

For the results reported above, we used animals from plates that were initially populated with five gravid adults and then incubated at 20 °C for three weeks. Under these conditions, the growing population depletes the bacterial food lawn after about five days, and thus these animals have starved for approximately 16 days. In contrast, the classical *C. elegans* freezing protocols call for freshly starved animals (Stiernagle, 2006).

Since starvation of *C. elegans* activates stress resistance pathways (Weinkove *et al*., 2006; Larance *et al*., 2015), and has been shown to upregulate endogenous trehalose production (Hibshman *et al*., 2017), we asked if starvation time may affect the rate of survival after desiccation.

We compared survival after drying in trehalose buffer of worms incubated for three weeks versus worms incubated for one week. We found that overall survival was significantly higher for worms incubated for three weeks (4.1 ± 0.6% versus 1.1 ± 0.4%, p = 0.009, Wilcoxon rank-sum test, 6 replicates with an average of 7,700 and 12,500 worms per replicate for three week and one week worms, respectively). However, we did not find a significant difference between survival rates of non-dauer (L1-L3) larval stages in the two groups (1.8 ± 0.5% for three week plates versus 0.95 ± 0.28% for one week plates, p = 0.22, Wilcoxon rank-sum test). While dauers were 7.6% of the total population on the three week plates, they made up only 0.5% of the population in the one week plates. Dauers from three week plates survived desiccation at an average rate of 32%; we did not measure the survival rate of dauers from one week plates due to their low numbers.

These results suggest that prolonged starvation increases survival after desiccation primarily by increasing the abundance of dauer larvae.

### The dauer defective mutant daf-16 does not survive desiccation

Some *C. elegans* mutants have a lower rate of survival than wild-type following freezing using conventional methods. One example is mutants for the forkhead box (FOXO) homologue DAF-16 (Hu *et al*., 2015). DAF-16 is involved in the response to cellular stressors, and *daf-16* loss-of-function mutants are dauer defective (cannot enter the dauer stage). To determine if *daf-16* mutants can be stored in desiccated form, we desiccated *daf-16(mu86)* mutants in our trehalose and glycerol supplemented buffers. Upon rehydration, no survivors were recovered from four replicates using trehalose buffer, and a single survivor was recovered from four replicates using glycerol buffer.

DAF-16 regulates production of endogenous trehalose and glycerol (Hibshman *et al*., 2017). However, following preconditioning on plates containing high NaCl, wild-type and *daf-16* mutants survive acute osmotic shock equally well (Lamitina and Strange, 2005). We hypothesized that preconditioning *daf-16* mutants on high NaCl plates might enable them to survive desiccation. To test this hypothesis, we dried and attempted to recover *daf-16* mutants raised on NGM plates containing an additional 200 mM NaCl. Out of three replicates for each protective agent, trehalose and glycerol, we did not recover a single survivor. This result shows that the dauer defective *daf-16* mutant does not survive desiccation, even when preconditioned to withstand osmotic shock.

## DISCUSSION

We have described a method for desiccating and freezing *C. elegans* in the presence of xeroprotective agents. Strains stored this way can survive cycles of warming and refreezing that accompany equipment failures or power outages. We provide a detailed version of our protocol in Supplementary File S2.

While *C. elegans* dauers (Erkut *et al*., 2011), and potentially other developmental stages (Ohba and Ishibashi, 1981), were previously shown to be capable of anhydrobiosis, we found that the addition of exogenous xeroprotective agents trehalose or glycerol dramatically improved survival rates. In the wild, *C. elegans* commonly inhabit decomposing fruit (Frézal and Félix, 2015). While most fruits do not contain trehalose, other sugars often found at high concentrations in fruits, including fructose, glucose, and sucrose, have been shown to act as cryoprotectants in some organisms (Storey and Storey, 1991) and may also function as xeroprotectants. Ethanol, a biproduct of sugar fermentation in rotting fruit (McKenzie and McKechnie, 1979), has been shown to increase *C. elegans* desiccation tolerance (Kaptan *et al*., 2020). Therefore it is possible that the ability to survive drying in the presence of exogenous sugar and other componants of *C. elegans’* natural surroundings may provide a selective advantage.

We demonstrated that dehydrated stocks can be frozen and thawed multiple times without significant loss of viability, and that they can be stored at warmer temperatures for shorter periods. The ability of desiccated stocks to retain viability at conventional refrigeration temperatures may be useful after a refrigeration failure.

A significant limitation of our method is that *daf-16* mutants, and perhaps others, do not survive drying and rehydration. As in conventional freezing, it is important to conduct a test thaw whenever preparing strains for storage.

Our attempts to precondition *daf-16* mutants by culturing them on high-salt plates did not improve survival after desiccation. It is possible that the expression of other DAF-16-regulated products, such as genes involved in the hypertonic stress response (Lamitina and Strange, 2005), is required in addition to exogenous cryoprotectants for enhanced survival after desiccation. It is also possible that the ability to form dauers is required for desiccation survival, and that other dauer defective mutants (Hu, 2007) would also fail to survive desiccation.

Our method may be useful for the exchange of strains between *C. elegans* laboratories. Strains are normally shipped on NGM plates, which are vulnerable to cold temperatures during transport and requires the sender to subculture worms before shipping. Shipping desiccated *C. elegans* would have the advantage of not requiring subculture of strains at either end, and would be impervious to cold, although desiccated worms remain vulnerable to heat.

There have been relatively few descriptions of cryopreservation of desiccated organisms. In freeze-drying (lyophilization), samples are frozen first, then dehydrated, then stored at low temperature or room temperature. While lyophilization is often done with microbes (Morgan *et al*., 2006), few animals or animal cells can be stored this way (Katkov *et al*., 2012). In contrast, our method involves *dry freezing, i.e*. first drying and then freezing the specimen. Reports of dry freezing as a practical storage method have been limited to the preservation of plant seeds (Pritchard, 2007) and seedlings (Sun, 1958). However, survival of freezing by dehydration seems to be relatively common in nature, particularly in soil-dwelling animals (Holmstrup, Bayley and Ramløv, 2002). Our method may suggest exogenous xeroprotectant-based dry freezing protocols in other systems.

## Supporting information

Supplemental Protocol File S2

## ACKNOWLEDGEMENTS

Strains were provided by the CGC, which is funded by NIH Office of Research Infrastructure Programs (P40 OD010440). C. F.-Y. was supported by the National Institutes of Health (R01 NS115995). D. M. R. was supported by National Institutes of Health (R01 NS107969 and R01 NS088432).

## Supplementary Materials

**Video File S1**: Time lapse video of dehydration, rehydration, and recovery of *C. elegans*. A 10 μL droplet of N2 worms in M9 buffer containing 15% trehalose was pipetted on a PDMS surface and dehydrated for 12 h. The dehydrated droplet was transferred to a seeded NGM plate for recovery.

**Protocol File S2**: Detailed protocol for dehydration, rehydration, and recovery of *C. elegans*.

