## Supplemental Protocol File S2 for "Dehydrated *Caenorhabditis elegans* stocks are resistant to multiple freeze-thaw cycles"

### Dry Freezing Protocol

#### Reagents, Materials, and Equipment

1. Desiccant (anhydrous calcium sulfate with cobalt chloride indicator, 8 mesh size, Drierite, W. A. Hammond, 20 g per strain)
2. M9 buffer (Stiernagle 2006)
3. Trehalose (e.g. Sigma T9449)
4. 15 mL conical centrifuge tube (1 per strain)
5. 1.8 mL cryogenic tubes (4 per strain)
6. 1.5 mL microcentrifuge tubes (2 per strain)
7. 6 cm diameter NGM OP50 plates (Stiernagle 2006) (2-3 per strain)
8. Sterile transfer pipette
9. Airtight box (approximately 28 X 19 X 8 cm, *e.g.* Rubbermaid 1991158)
10. Rack for cryotubes (must fit in the airtight box)
11. Cryostorage box
12. Platinum worm pick (must reach bottom of cryotube)
13. P200 and P1000 micropipettes
14. Centrifuge
15. Microbalance
16. Vortexer
17. Stereo microscope

#### Dehydration and freezing procedure

1. Pick five gravid adults onto each of 2-3 standard 6 cm OP50 plates, and incubate at 20 °C for three weeks. After three weeks, ensure that plates are free of contamination, that the animals have depleted their bacteria food, and that some dauer larvae are present.

2. Mix 30% w/v trehalose in M9 (*e.g.* 150 µg trehalose in 500 µL M9) in a 1.5 mL microcentrifuge tube and set aside.

3. Use a sterile transfer pipette (see Note 1) and several mL of M9 to wash the worms into a 15 mL conical centrifuge tube. Add M9 to a final volume of 10 mL.

4. Pellet worms by centrifuging for 5 min at 700 rcf. Use a sterile transfer pipette to remove the supernatant. Add fresh M9 to 10 mL, pellet worms, and remove supernatant again, leaving < 100 µL of M9 with the pelleted worms.

5. Use a sterile transfer pipette to transfer the pellet to a 1.5 mL microcentrifuge tube. Add M9 to 125 µL using a microbalance to aid with measurement (see Note 2).

6. Add 125 µL of 30% trehalose in M9, and mix by gentle vortexing or flicking the tube. This is the "worm mixture".

7. Pipette 50 µL of worm mixture into each of the four cryotubes.

8. Place the *uncapped* cryotubes upright into a tube rack or other support inside the drying box

9. Scatter 20 g of fresh desiccant onto the bottom of the drying box.

10. Seal the lid and leave the drying box at RT for 48 h.

11. Open the drying box. Test one tube for dryness by using a stereo microscope to observe the droplet of worm mixture while touching it with a sterile worm pick. A properly dried drop should be hard around the edges, but scratch or dent slightly in the center (see Notes 3 and 4).

12. Cap the cryotubes and place them in a cryostorage box at the desired ultracold storage temperature.

#### Recovery procedure

1. Thaw tube for 10 min at RT.

2. Add 50 µL of M9, and incubate 10 min at RT.

3. Gently pipette the rehydrated worms up and down to resuspend. Pipette the worm suspension onto an NGM-OP50 plate.

4. Check for survivors the following day.

#### Notes

1. The same sterile transfer pipette can be reused in steps 3, 4, and 5 provided that it is kept in its sterile wrapper when not in use and only used for one strain.
2. A microbalance can help with this step. Weigh and tare the empty 1.5 mL microcentrifuge tube and assume an approximate density of 1 g/mL for the M9-worm mixture.
3. Our observations suggest that for optimal worm survival a droplet should be neither too wet nor too dry. Factors such as droplet volume, air humidity, and temperature can affect droplet dryness. To increase droplet dryness, use more desiccant; to reduce dryness, use less desiccant.
4. Using a sealed box and the ratio suggested, we observed little or no color change in the desiccant. Excessive color change could indicate that the drying box is not well sealed.
